# Force transduction creates long-ranged coupling in ribosomes stalled by arrest peptides

**DOI:** 10.1101/2020.10.16.342899

**Authors:** Matthew H Zimmer, Michiel JM Niesen, Thomas F Miller

**Affiliations:** California Institute of Technology; Massachusetts Institute of Technology

## Abstract

Force-sensitive arrest peptides regulate protein biosynthesis by stalling the ribosome as they are translated. Synthesis can be resumed when the nascent arrest peptide experiences a pulling force of sufficient magnitude to break the stall. Efficient stalling is dependent on the specific identity of a large number of amino acids, including amino acids which are tens of angstroms away from the peptidyl transferase center (PTC). The mechanism of force-induced restart and the role of these essential amino acids far from the PTC is currently unknown. We use hundreds of independent molecular dynamics trajectories spanning over 120 μs in combination with kinetic analysis to characterize the barriers along the force-induced restarting pathway for the arrest peptide SecM. We find that the essential amino acids far from the PTC play a major role in controlling the transduction of applied force. In successive states along the stall-breaking pathway, the applied force propagates up the nascent chain until it reaches the C-terminus of SecM and the PTC, inducing conformational changes that allow for restart of translation. A similar mechanism of force propagation through multiple states is observed in the VemP stall-breaking pathway, but secondary structure in VemP allows for heterogeneity in the order of transitions through intermediate states. Results from both arrest peptides explain how residues that are tens of angstroms away from the catalytic center of the ribosome impact stalling efficiency by mediating the response to an applied force and shielding the amino acids responsible for maintaining the stalled state of the PTC.

**Significance Statement:** As nascent proteins are synthesized by the ribosome, their interactions with the environment can create pulling forces on the nascent protein that can be transmitted to the ribosome’s catalytic center. These forces can affect the rate and even the outcome of translation. We use simulations to characterize the pathway of force transduction along arrest peptides and discover how secondary structure in the nascent protein and its interactions with the ribosome exit tunnel impede force propagation. This explains how amino acids in arrest peptides that are tens of angstroms away from the ribosome’s catalytic center contribute to stalling, and, more broadly, suggests how structural features in the nascent protein dictate the ribosome’s ability to functionally respond to its environment.

## Introduction

Regulation of protein expression levels is an essential part of maintaining cellular homeostasis. While this is most commonly performed at the nucleotide level by controlling rates of transcription initiation or ribosome binding, regulation at the translational level allows for immediate control of protein levels [1]. One method of translation-level regulation utilizes arrest peptides, which are short amino acids sequences that induce a stall in ribosomal translation when synthesized, until an external signal breaks the stall. [2]. Force-sensitive arrest peptides (FSAPs) are a class of arrest peptides that allow for the restart of translation with a force-dependent rate [3]. Physiologically, these peptides provide a force sensor associated with the integration of nascent proteins into the membrane or translocation across the membrane. For example, the *E. coli* arrest peptide SecM is able to regulate expression of the translocase SecA because restart of SecM-stalled translation is dependent on the pulling force exerted by SecA on the nascent peptide [4]. Due to the sensitivity of SecM stall breaking to external forces, SecM has also been used as a biophysical tool to measure forces *in vivo* [5, 6, 7]. Understanding the mechanisms by which FSAPs stall the ribosome and respond to forces will not only shed light on an essential mode of regulation in biology, but also provide insights for the design of mutants that respond to forces with greater dynamic range and sensitivity, thereby enabling wider application of FSAPs as *in vivo* force sensors [8, 9].

Considerable progress has been made in understanding the molecular mechanism behind FSAP stalling. SecM was the first FSAP identified and remains one of the best characterized. The requisite sequence in SecM for stalling has been narrowed down to a stretch of seventeen amino acids, of which nine amino acids are essential for efficient stalling[10, 11]. Cryo-EM structures have identified several SecM stalled states with different ribosome conformations and tRNA occupancies [12, 13]. A common feature of the different stalled conformations is that the SecM sequence alters the geometry of the ribosome in such away as to stabilize the uninduced state, blocking the addition of the next amino acid. The importance of particular residues is also made clear by these structures; for example, the essential C-terminal R163 interacts with several neighboring nucleotides to distort the peptidyl transferase center (PTC) geometry [14]. However, other amino acids essential for efficient stalling, such as F150 and W155, are more than ten angstroms away from the PTC and have no obvious role in altering the conformation of the ribosome. Although structural data show the essential residues far from the PTC are interacting with ribosomal proteins and RNA in the exit tunnel, there are no large scale rearrangements in the ribosome that could lead to propagation of this signal through the ribosome [15, 12, 16, 17]. Instead, these residues are suggested to increasing stalling by precisely arranging the conformation of C-terminal portion of the nascent chain or by controlling the degree of compaction in exit tunnel [9, 18]. The molecular details of how these essential C-terminal amino acids induce this arrangement have not yet been uncovered.

More recently, the structures of ribosomes stalled by VemP and MifM, two other FSAPs, have been solved [19, 17]. These peptides share many of the same features as SecM, namely the stabilization of the uninduced state of the ribosome and the presence of many essential amino acids positioned far down the exit tunnel. Despite these similarities, there are also significant differences between these more recently characterized peptides and SecM, most notably the secondary structure and compaction of the nascent chain in VemP and the strict species specificity of MifM.

While the available structures provide insight into how arrest peptides stall translation, the mechanism of force-dependent restart is less well understood. In the current work, a molecular picture of the stall-breaking processes is revealed by microsecond-timescale molecular dynamics (MD) simulations of the SecM and VemP FSAPs with an applied force on the N-terminus, yielding over 150 independent stall-breaking events. These simulations uncover a conserved multi-step pathway in which pulling forces are unable to disrupt the conformational around the PTC until key interactions formed by the N-terminal regions of the arrest peptides are broken. We identify the role by which essential amino acids far from the PTC affect force-dependent restarting and a mutation strategy to produce FSAPs with altered force sensitivity. Comparison between the stall-breaking pathways of SecM and VemP highlights the different effects on force transduction of interactions between the nascent peptide and the ribosome as compared to intra-chain interactions in nascent peptide. Beyond FSAPs, co-translational forces have been shown to be able to influence the rates and even outcomes of translation [20, 21]; thus, a better understanding of the factors that control force propagation in the exit tunnel also provides valuable insight into how ribosomes are able to sense and respond to their environment via the nascent protein.

## Results

### Pulling-force trajectories

The force-induced restart of stalled ribosomes is a non-equilibrium biomolecular process that occurs on the timescale of seconds to minutes. Microsecond-timescale molecular dynamics with an applied force on the nascent protein provides insight into these non-equilibrium effects and allows for the direct visualization of the motions and forces of the nascent chain throughout the stall-breaking pathway. Simulations are initialized using cryo-EM structures of SecM and VemP stalled ribsomes. SecM stalls the ribosome at several states during synthesis and several of these states have been characterized structurally [22, 12, 13]. Simulations of SecM were started from the structure corresponding to the earliest point in translation (PDB: 3JBU), in which Pro-tRNA has not yet bound to the ribosome (Fig. 1A). To reduce computational cost, the ribosomes in both SecM and VemP structures are truncated beyond a 23 Å radius around the nascent chain, with an outer shell of 3 Å restrained to preserved the structure of the ribosome (Fig. 1B, Methods). Following truncation, the starting structures were relaxed using 1 μs of equilibrium MD.

**Figure 1:**
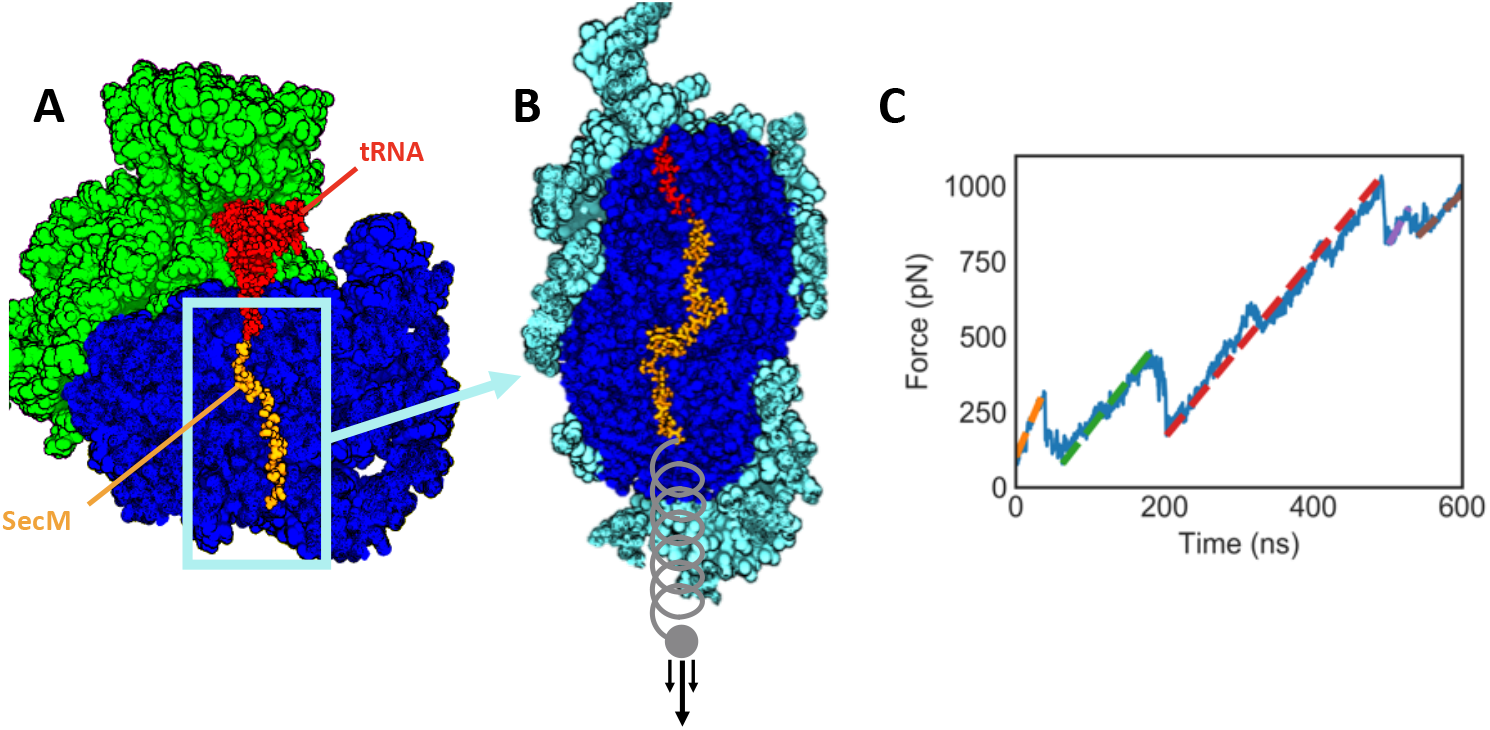
Description of the simulation system and typical output. A: Cryo-EM structure of a SecM stalled ribosome (PDB: 3JBU). Only the region of the ribosome surrounding the nascent chain is included in our simulations. B: The simulated part of the ribosome, with atoms shown in cyan harmonically restrained. A ramping force is applied to the N-terminus of the nascent chain. C: The result of a SecM pulling trajectories. Several regions of linear extension can be identified (colored dashed lines), corresponding to different stable conformations of the nascent chain. Note that, for visual clarity, the structures in A and B do not show atoms that are nearer to the viewer than the nascent chain.

To simulate force-induced stall breaking, we apply a force on the nascent chain by placing the N-terminal alpha carbon in a harmonic potential, with the minimum energy position of the harmonic potential moving out of the exit tunnel at constant velocity. This protocol mimics the commonly used force-ramping protocol in optical tweezer experiments [3, 23, 24]. To observe stall-breaking on a timescale accessible to simulation, it is necessary to pull with a faster force-ramping profile than is done experimentally. We run simulations over three orders of magnitude in force loading rates (0.5, 5, and 50 pN/ns). These loading rates lead to rupture forces between 100 to 1000 pN and timescales from 50 ns to 5 μs. Although this is faster than the presumed experimental timescale of seconds [3, 22], the wide range of pulling forces applied allows us to evaluate the consistency of our trajectories with varying simulation timescale. The analysis of over one hundred independent trajectories allows for more detailed and statistically robust analysis than previous simulations of stalling peptide restart [14, 25]. The large number of long trajectories (over 120 μs total) was enabled by the use of the Anton2 computer [26].

### Force propagation through stall-breaking intermediates

We first consider force-pulling simulations involving the SecM sequence, as SecM is the best characterized FSAP and has seen the most use in force-measurement experiments, enabling us to validate our methodology and make readily testable predictions. The force profiles indicate that several distinct conformational changes occur within a typical trajectory (Fig. 1C). The changes are evidenced by the sharp drops in the force profile, corresponding to moments in which the nascent chain extends along the direction of the applied force. Regions of the force profile in between these drops with linearly ramping force correspond to metastable intermediate states along the stall-breaking pathway. To identify states that are consistently observed across the 5 pN/ns trajectories, the linear rate of force increase is subtracted from each force profile (Fig. 2A, Methods). This allows for direct comparison between force profiles even though the times at which conformational changes occur are stochastic [27].

**Figure 2:**
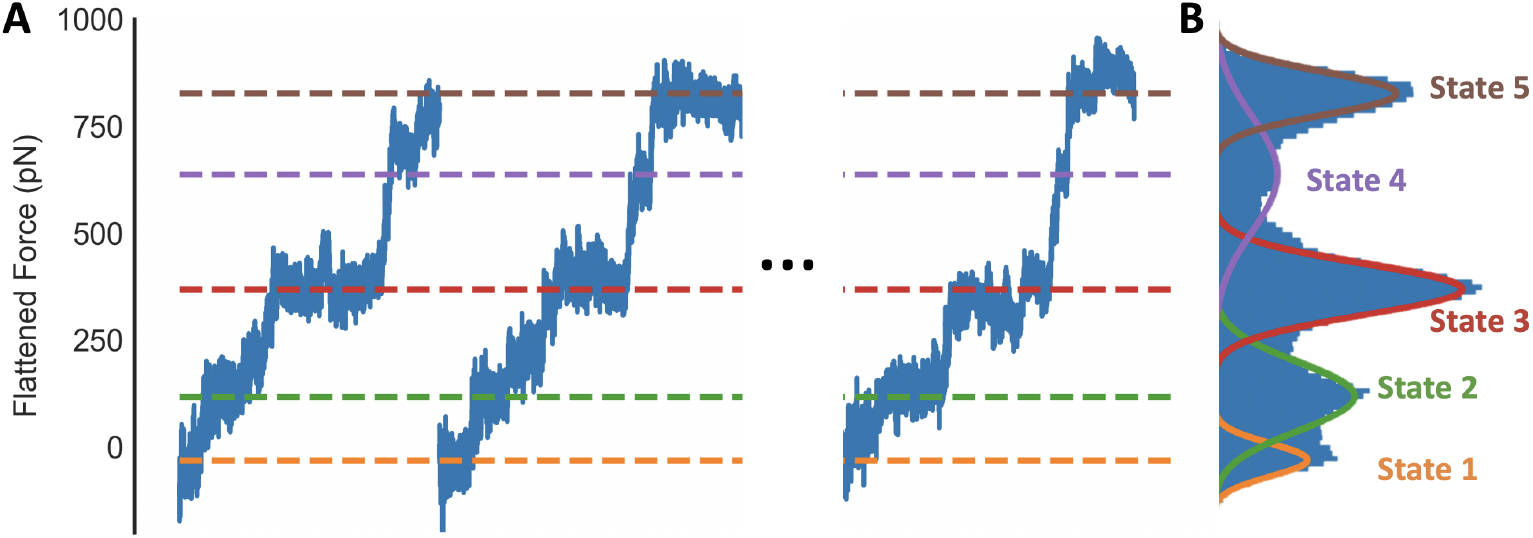
Identifying states based on force profiles A: Force profiles from three representative simulations of SecM after a force ramp of 5 pN/ns has been subtracted. Dashed lines indicate the peaks of the Gaussians which have been fit to the data in panel B. B: Histogram of the flattened force profiles of all thirty trajectories run with a force loading rate of 5 pN/ns.

Combining the results of the ensemble of pulling-force trajectories reveals a distribution of applied forces with five peaks, with each peak corresponding to a conformational state that is robustly observed across the trajectories (Fig. 2B). These data are then fit by the sum of five Gaussian distributions. Each frame of the trajectories is assigned to one of the five states based on the force at that frame. Each state may contain sub-states that give rise to small drops in the force profile within a given state (e.g. in Fig. 1C); however, these sub-states are not consistent across different trajectories. Since these sub-state transitions are not conserved in the stall-breaking pathway and their infrequent occurrence makes robust analysis difficult, they were not further characterized. Repeating the above analysis on simulations conducted with a faster, 50 pN/ns loading rate reveals a distributions of external forces that are also well-fit assuming a five state model (Fig. S1).

With thousands of configurational samples per state, the molecular motions that govern the transitions between the observed states can be characterized. Measuring the changes in dihedral angles between consecutive states on an per-amino acid basis (Fig. 3, SI Movie) reveals that the conformational changes between states are driven by rotations of only a few adjacent amino acids per transition. For all of the amino acids that undergo changes in dihedral angle, the rotation observed involves the straightening of the ψ dihedral angle from a bent conformation to a 0 degree planar conformation. This extends the nascent chain in the direction of the applied force. The set of residues that rotate in each transition progresses toward the C-terminus of nascent chain with each successive transition, from Ser151 rotating in state 1, Ser157 and Gln158 in state 2, Gly161 in state 3, and finally to Arg163 and Ala164 rotating at the C-terminus in state 4. The molecular descriptions of the states and the transitions between them are almost identical when simulations are performed with a 10x greater force ramping rate (Fig. 3). The same analysis performed on trajectories run with the slower ramping rate of 0.5pN/ns identified nearly identical conformational changes for each state transition, albeit with some overlap between the peaks, likely due to the difficulty in assigning states with only five trajectories (Fig. S2). The rotation of the specific amino acids listed above defines a robust molecular pathway that describes that conformational changes SecM undergoes when subjected to an applied force on the N-terminus.

**Figure 3:**
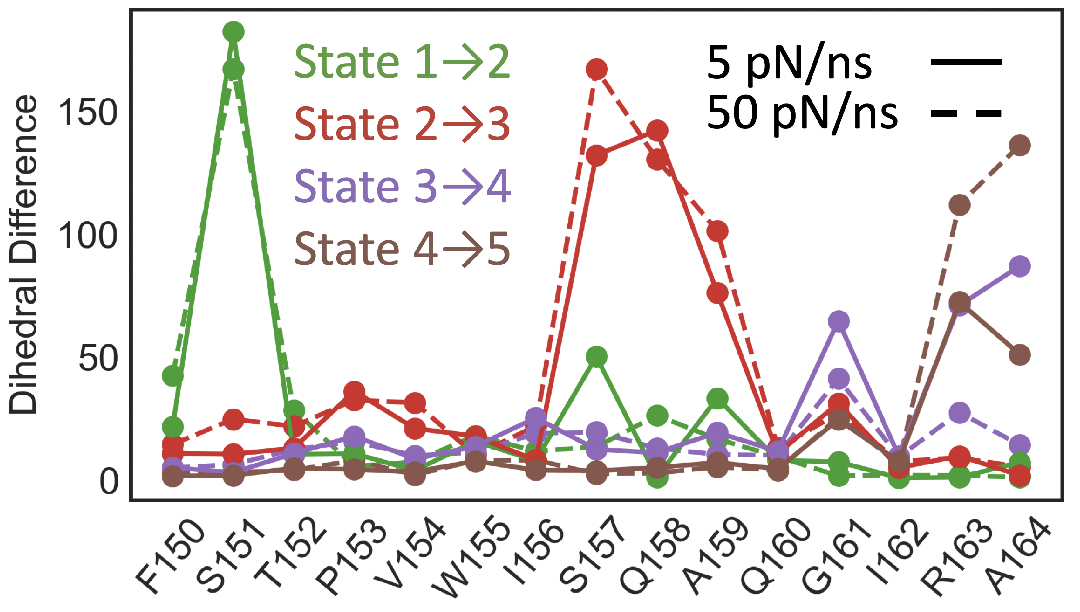
Conformational changes are quantified for each amino acid in terms of the distance of the ϕ and ψ dihedral angles between two states. The key differences between each state can be described by rotations in only a few amino acids per state.

### Connecting peptide conformation to restart of translation

We now identify the intermediate state during force propagation that most likely coincides with the restart of ribosomal translation. The five amino acids at the C-terminus of SecM are all essential to its stalling behavior, as well as being the closest to the site of new bond formation [9]. It is therefore expected that only conformations in which the essential C-terminal residues have been affected by the force could allow restart of synthesis. The most dramatic movement of these amino acids occurs during the transition from states 3 to 4 when Arg163 and Ala164 straighten (Fig. 3). However, this transition is unlikely to occur under physiological conditions as it coincides with an unphysical extension of the tRNA; characterizing the movement of the tRNA nucleotide which is covalently bonded to the nascent protein reveals that the tRNA maintains a stable position until the transition to state 4, at which point it is pulled several angstroms into the exit tunnel (Fig. 4A). This extension is enabled by the breaking of the stacking interactions between the nucleobases in the tRNA (Fig. S3) in a manner which, to our knowledge, has not been observed *in vivo*. To further evaluate the possible relevance of the 3-to-4 transition, we estimated the height of the free energy barrier by fitting the observed distribution of state transition times to a probabilistic model of force-induced barrier crossing proposed by Bullerjahn et al (Fig. S7, S8, Table S1) [28]. This leads to an estimate for the barrier height of 120kT (90% CI: 110-160 kT), which could not be crossed on biologically relevant timescale without forces far stronger than what can be exerted by SecA, estimated to be around 10 pN [29]. The analyses of the conformational changes and the energetics of the state 3 to 4 transition both indicate that state 4 is physiologically inaccessible and therefore that restart must occur in state 3 or before.

**Figure 4:**
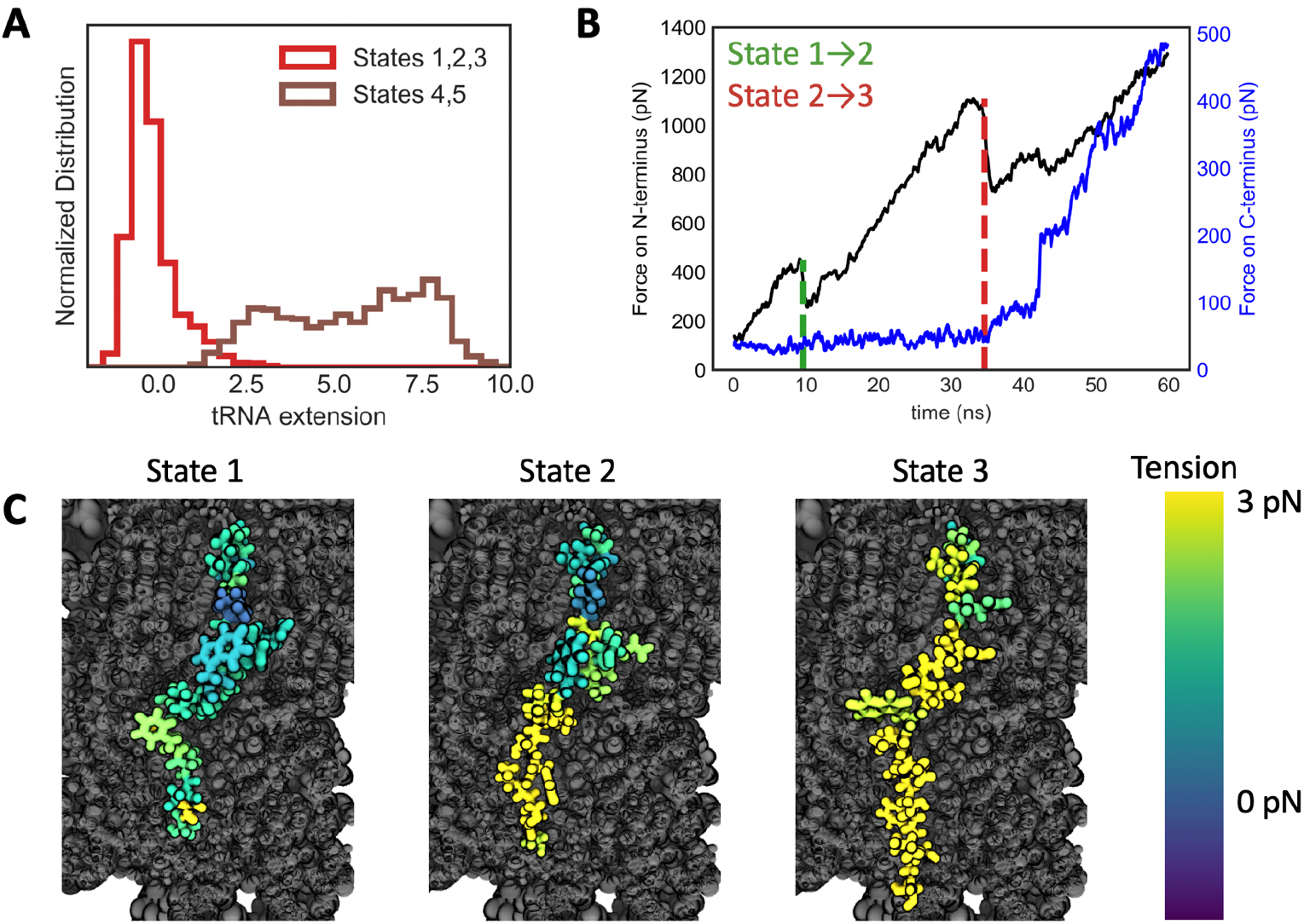
Identifying which states could correspond to restart of synthesis A: Distributions of the extension of the 3’ hydroxyl group of the tRNA nucleotide which is bonded to the nascent chain beyond its position during equilibration. Extension is measured in the direction of pulling. States 4 and 5 are unphysically extended. B: Simulations of SecM were run with the force on the N-terminus ramped at 50 pN/ns as previously, but now with the C-terminal Arg163 was fixed by a harmonic restraint. The pulling force is plotted in black and the amount of force that is acting on Arg163 is plotted in blue. The force only increases above its basal level until after the transition to state 3. The transitions between states 1 to 2 and 2 to 3 were determined by the frame in which residues Ser151 and Ser157 respectively straighten. C: Characteristic conformations of each state are shown, colored by the magnitude of the tension along the nascent chain relative to the tension in the equilibration trajectory. Tension only reaches the C-terminal amino acids once state 3 is reached.

For restart to occur, the conformation of the amino acids at the PTC must be disrupted by the applied force. This occurs only in state 3 and later (Fig. 3), suggesting that the transition from state 2 to 3 is necessary for restart of synthesis to occur. To precisely determine when the amino acids at the PTC experience pulling forces, simulations are run with a harmonic restraint on the alpha carbon of the essential Arg163 (Fig. 4B). The displacement of the restrained alpha carbon reports on the force that atom experiences. Until SecM enters state 3, no significant forces are felt by Arg163 and restart of translation cannot occur. This held true across several different simulation protocols, including with two different loading rates and when the restraint was placed on Gly165 rather than Arg163 (Fig. S4). The inhibited force transmission in SecM contrasts with previously published results that measured force transmission in a non-stalling peptide and found no significant difference between the forces at the N- and C-termini [20].

To further confirm the mechanistic significance of the transition from state 2 to state 3, the tension along the backbone of the nascent chain is measured (Methods). Tension in the nascent chain around the PTC only increases after state 3 is reached (Fig. 4C). Analysis of the computed tensions as a function of position along the nascent chain leads to the same conclusions as those reached above, while having the advantage of not requiring an artificial restraint and providing measurements from all SecM amino acids, instead of just the restrained atom. However, the tension fluctuates dramatically with time and meaningful comparisons can only be made after averaging over all the frames in a state.

Both metrics of tracking the intra-chain forces during a pulling trajectory indicate that overcoming the first barrier and entering the second state could not lead to a restart of translation. Only once the third state is entered does the force reach the C-terminus and enable disruption of the interactions around the PTC that hold the stall in place. Additionally, our estimate of the free energy barrier between states 2 and 3 is 27 kT (90% CI: 21-76 kT), which makes this transition the last energetically accessible step. The 4 kT (90% CI: 3-8 kT) barrier between states 1 and 2 can easily be overcome by thermal motion, whereas the 120 kT (90% CI: 110-160 kT) barrier between steps 3 and 4 is physiologically insurmountable, requiring the disruption of tRNA stacking. Taken together, this indicates that arrival at state 3 is both necessary and sufficient for stall-breaking.

### Force transduction in mutant arrest peptides

The extent to which mutations in SecM affect the response to an applied force is poorly characterized. Previous studies of stalling by SecM mutants were performed in the absence of the translocation machinery that physiologically exerts the forces necessary to relieve the stall [10, 9, 18]. These experiments identify which amino acids are essential for stalling, but do not indicate which mutants alter force sensitivity. We thus employ MD simulations to explore how mutants alter the barriers to force propagation along the nascent chain. Several amino acids in SecM were mutated to alanine, equilibrated for 250ns, then re-simulated with a force loading rate of 50 pN/ns. From these trajectories, the work required to pull the mutant into the restart-competent state 3 is calculated. To focus on the effects due to force propagation rather than the conformational changes that may occur upon mutation, mutations were made only to amino acids more than 10 Åaway from the PTC. Five mutants were simulated, three that were found experimentally to reduce stalling efficiency in the absence of an applied force (F150A, W155A, I156) and two that were not observed to have an effect on stalling (L149A, T152A). The three mutants that required the least amount of work to restart were the same three that were found to experimentally reduce stalling efficiency (Fig. 5B) [10]. This indicates that the same amino acids that are necessary to arrange the upper portion of the nascent chain into a stalling conformation [18] also play a significant role in mediating the response to a pulling force.

**Figure 5:**
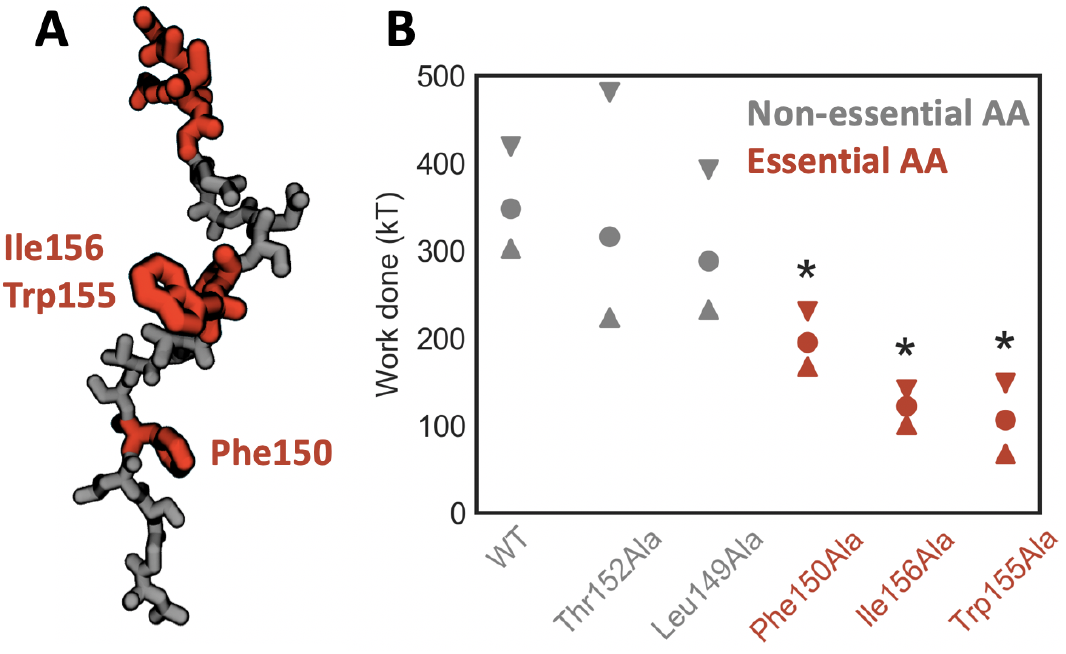
Exploring the role of the essential amino acids in the restart process A: Snapshot of SecM with the amino acids whose identity is essential to effective stalling colored in red. B: Mean work done to reach the restart-competent state 3. The mutants colored grey are those which were experimentally found to have little affect on stalling, while the red mutants led to a significant reduction in stalling. Error bars indicate 95% confidence intervals on the mean. Stars indicate mutants for which the mean of the work done was significantly different from the WT work according to a t-test with unequal variances and a threshold of *p* < 0.01

These findings have broad implications for the process of force-induced restarting of stalled peptides. Firstly, it provides an explanation for why the specific identity of amino acids is essential even when they are tens of angstroms away from the PTC, as the force is unable to propagate beyond these amino acids until specific interactions with the ribosomal exit tunnel are broken (Fig. 4A). The interactions formed by these residues should also be capable of blocking or damping fluctuations on the chain, which provides further connection to mutagenesis studies that investigate stalling behavior at zero applied force [10, 11, 18]. This mechanism also suggests that mutating these essential amino acids lower in the chain would be the most effective strategy for tuning the force response properties of SecM. Mutations of the essential residues at the C-terminus, such as Arg163, would be more likely to abrogate the stall entirely rather than change its sensitivity to an applied force. Additional variants of SecM with altered response to force would aid the use of FSAPs as in vivo force sensors, since the diversity biological processes that arrest peptides can report on is limited by the dynamic range their force induced restarting[30].

### Alternative stall sequences

Simulations of SecM suggest a mechanism in which interactions between the nascent chain and the ribosome inhibit force propagation to the PTC and prevent restart of translation until those interactions are disrupted. To evaluate the generality of this mechanism, a second force-sensitive arrest peptide was investigated, VemP. As with SecM, force-sensitive stalling of VemP regulates the expression of protein translocation components[31]. However, a recent structure of VemP stalled in the ribosome exit tunnel showed that VemP adopts a conformation in the exit tunnel highly dissimilar to SecM [17] (Fig. 6A). Unlike SecM, the VemP nascent chain is highly compacted with clear secondary structure; two helical regions in the upper and lower sections of the peptide. The upper helix is principally responsible for inducing the stall, with several amino acids interacting with ribosomal nucleotides in such a way as to block peptide bond formation. It has been proposed that an force applied to the nascent chain could unravel the upper helix, thereby disrupting these interactions with the ribosome and enabling the restart of translation [17]. Due to the very different stalled conformation of VemP, it provides a stringent test of the generality of the mechanism uncovered for the force induced-restarting of SecM.

**Figure 6:**
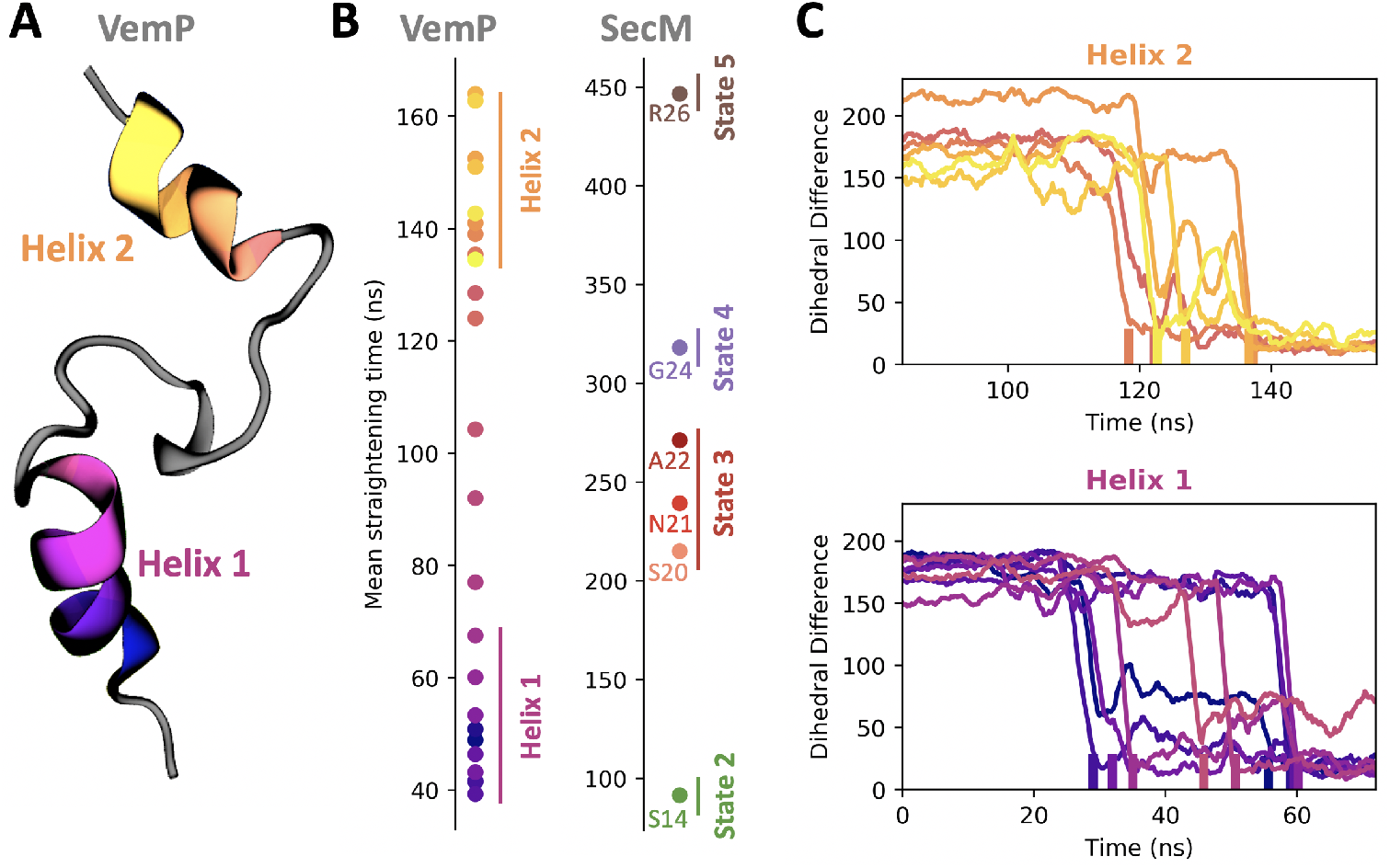
Force-induced conformational changes in VemP. A: Structure of the stalled VemP nascent chain, with the two helical regions colored. B: Straightening time for different amino acids in VemP and SecM averaged over 30 trajectories, using a force loading rate of 35 pN/s. C: Time series of the deviation of the dihedral angles in helices 1 and 2 from the fully extended conformation, taken from a representative trajectory. The amino acids within a helix do not straighten in consecutive order.

Following the same protocol used to study SecM, simulations of VemP were prepared from its cryo-EM structure (PDB: 5NWY), with the ribosome cut-off and restrained beyond a 23 Å radius. Force was applied on the N-terminus along the direction of the exit tunnel with a force-loading rate of 35 pN/s. Due to the high degree of compaction in VemP, if the nascent chain were fully extended from its starting configuration it would extend well outside the ribosome exit tunnel. To reduce the total simulation volume, a pulling trajectory is divided into three parts and the fully extended portion of the nascent chain is removed at the end of each third (Methods). This relies on the assumption that the fully extended nascent chain will not inhibit the propagation of the applied force.

Overall, the N-terminal helix unravels strictly before the C-terminal helix, as measured by the average time at which the dihedral angles of each residue straighten (Fig. 6B). This suggests that, as with SecM, interactions formed further from the PTC must be broken before force can propagate to residues nearer the PTC. Tracking tension throughout the simulations also indicates that the tension in the C-terminal helix near the PTC does not increase until the N-terminal helix is unraveled (Fig S5).

However, the presence of secondary structure in VemP also leads to notable differences in the mechanism of restart. One such difference is that the straightening of amino acids *within* a helix does not necessarily occur consecutively. This can be seen both in the straightening times of the amino acids averaged over all trajectories (Fig. 6B) and individual trajectories (Fig. 6C). In Fig. 6, the amino acids and the markers for their transitions are colored from darker to lighter from the N-terminus to the C-terminus. If the order of straightening was strictly consecutive, the ordering of colors in Fig. 6 B and C would become monotonically lighter with time. This is evidently not the case within both helices 1 and 2. In fact, not only is the order of unravelling of different turns of a helix not consecutive, it is also not conserved across independent trajectories simulated with the same protocol (Fi. S6).

The underlying reason for the difference in ordering between SecM and VemP arises from the fact that the interactions found in the SecM simulations are between the nascent chain and the ribosome, whereas the interactions within a VemP helix are between different VemP amino acids. Interactions with the nascent chain and ribosome hold the nascent chain in place, relative to the exit tunnel. Intra-chain interactions in a helix still allow the helix to move as whole in response to a force, which allows force to propagate through the helix. This means that interactions within and directly C-terminal of a helix break in order of their free energy barriers. For example, if there is a weak interaction between the nascent chain and the exit tunnel C-terminal of a helix, then that weak interaction can be broken by an applied force before the alpha helix is disrupted. Conversely, if the exit tunnel-nascent chain interaction is stronger than the intra-helical interactions, then the alpha helix will be disrupted first. The similar free energy barriers for unfolding different turns of the same helix is likely why heterogeneity is observed in the breaking order within a helix between trajectories. This difference between inter- and intra-chain interactions can also explain why the middle portion of VemP which lacks secondary structure straightens in consecutive order.

Due to the greater cost of simulations of the longer VemP peptide, there are still several open questions regarding how its secondary structure responds to a pulling force. For instance, we were not able to investigate how the order of unraveling within a helix depends on an applied force, although we hypothesize that the heterogeneity would only increase at lower forces as thermal motion becomes more important [32]. The lack of trajectories at longer timescales also prevents estimation of free energy barriers, as was done with SecM. While challenging, this could be a particularly fruitful direction for future simulations as the barrier height determines whether nascent chain-ribosome or intra-chain interactions will break first. Additionally, although it was not observed in these trajectories, steric interactions between an alpha helix and a constricted region in the exit tunnel may force consecutive unraveling of the alpha helix as it needs to pass through the constriction zone. Despite these open questions, simulations of VemP revealed clear similarities and differences as compared with SecM. VemP exhibits non-consecutive breaking of interactions within a helix, while the interactions in SecM are disrupted consecutively from the N-terminus to the C-terminus. Importantly, both stalling peptides still require N-terminal interactions between the nascent chain and the exit tunnel to be broken before the critical conformation stalling the PTC can be disrupted.

## Conclusion

The results presented here reveal a mechanism for force-induced restart of ribosomal translation in which co-translational force propagation is governed by both interactions between the nascent chain and the ribosome and intra-nascent chain interactions. Simulations identified a pathway of sequential conformational changes in SecM that are required in order for a force applied at the N-terminal end of SecM to propagate to the PTC. This pathway was found to be consistent over two orders of magnitude of force ramping rate, and mutations that were experimentally found to reduce the stalling efficacy of SecM in the absence of force also reduced the work required to progress through the various conformational changes. Analysis of the tension in the VemP nascent chain also showed that the N-terminal interactions in helix 1 must be unravelled before helix 2 could be unfolded by the force, despite considerable heterogeneity in the unfolding pathway within individual helices. These results highlight a previously underappreciated distinction between amino acids in FSAPs which induce stalling through specific interactions with the ribosome around the PTC and those which stabilize the conformation of the nascent chain and shield the residues at the PTC from external forces. We believe the latter interactions should be the principal target of mutagenesis aimed to alter the force sensitivity of FSAPs, since mutations to amino acids around the PTC are likely to reduce stalling efficiency even in the absence of an applied force

Understanding co-translational force propagation is not only essential for understanding the restarting of FSAPs but also other co-translation processes that can be controlled by force, such as translation rate or frameshifting [20, 21] Comparison between the pulling force simulations of SecM and VemP emphasizes the role of interactions between the nascent chain and the exit tunnel which block further propagation of force up the nascent chain. This is contrasted with intra-chain interactions such as in alpha helices which can move as a unit and therefore do not inhibit the propagation of force and allow breaking of interactions in non-consecutive order. The force-induced restarting pathways of SecM and VemP both highlight the need to consider potential interactions between the nascent chain and the ribosome when estimating the co-translational forces experienced by the ribosome [33, 34]. Fortunately, characterization and understanding of interactions between a nascent peptide and the exit tunnel are becoming increasingly available thanks to the rapid expansion of cryo-EM structural data. Future simulations will provide an essential connection between the study of nascent chain behavior in the exit tunnel [35, 36, 37] and the role of forces in influencing cotranslational behavior[34, 21].

## Methodology

### Simulation set-up

Simulations of SecM were initialized from the 3.6 Å resolution PDB structure 3JBU [12]. System preparation was performed using Schröodinger Maestro. First, the covalent peptidyl-tRNA bond was manually restored. The structure was then truncated by removing amino acids with any heavy atoms 24 Å beyond the region of interested, defined as the SecM nascent chain and first three nucleotides of the P-site tRNA. This region was chosen to sufficiently large to minimize any possible influence of the truncated region, as they lie beyond the cut-off distance for the non-electrostatic interactions of the MD forcefield. Any amino acid or nucleotide monomers created by the truncation were removed. The SecM nascent chain was cut off at Leu149, and a SerGlySer linker was appended to the N-terminus. Hydrogens were modeled in, and N- and C-termini newly created by the truncation were capped with acetyl and N-methyl amide groups, respectively. Minimization was performed to alleviate clashes formed by the newly added capping groups. The structure was then solvated with TIP3P water and neutralized with Mg ^2+^. Solvation filled the orthorhombic periodic simulation box, which was set to extend 10 Å beyond the truncated ribosome. The ribosome was oriented to minimize the volume of the box. To retain the structure of the ribosome despite the truncation, all non-solvent heavy atoms 20 Å away from the region of interest were restrained with 1 kcal/mol/Å harmonic restraints. Initial simulation was run for 30 ns using the AMBER99SB-ILDN forcefield [38] and the Desmond simulation package [39]. The same steps were used to prepare the VemP structure from the 2.9 Å resolution PDB structure 5NWY, with no truncation of the nascent chain [17]. Equilibration was then performed on Anton2. SecM was equilibrated for 1000 ns and VemP for 500 ns. Simulations of mutant SecM were prepared starting from the SecM structure after equilibration. After mutations were introduced, each structure was re-equilibrated for 250 ns.

### Force-pulling protocol

The pulling direction was determined by identifying the vector that extends from the N-terminal alpha carbon of SecM or VemP and maximizes the distance between any point along the vector and the atoms of the ribosome. The optimal vector passes through the exit tunnel and maintains a distance of greater than 6 Å from any ribosomal atoms at all times.

Pulling is performed by applying a 1 kcal/mol/Å harmonic potential to the N-terminal alpha carbon that constrains the the distance between the alpha carbon and a point 30 Å away from the alpha carbon in the direction of pulling vector. Initially, this distance is set to the equilibrium distance of 30 Å, such that there is no force applied to the alpha carbon. This distance is decreased linearly with time, applying increasing force to the N-terminal alpha carbon. We explore different force loading rates of 0.5, 5, and 50 pN/ns. Simulations for were run 6.5 μ, 650 ns, and 78 ns respectively. These times correspond to halting the simulation when the minimum of the harmonic potential used to apply the pulling force was 45 Å away from initial position of the SecM N-terminus. For the fastest force loading rate of 50 pN/ns, this was extended to 55 Å to ensure all conformational transitions were observed. 30 independent simulations were performed at the force loading rates of 5 and 50 pN/s. Due to the longer time required, only 5 simulations were run with a force loading rate of 0.5 pN/ns. For all simulations, frames are recorded every 1.5 ns.

Simulations to measure the force at the C-terminus of SecM were performed as described above, but with the addition of a 1 kcal/mol/Å harmonic restraint on the alpha carbon of Arg163 or Gly165. The minimum of the harmonic potential was centered on the alpha carbon’s position in the cryo-EM structure. 25 independent trajectories were run with a loading rate of 50 pN/s and C-terminal restraint on Arg163, 10 trajectories with a loading rate of 5 pN/s and C-terminal restraint on Arg163, and 10 trajectories with a loading rate of 50 pN/s and C-terminal restraint on Gly165.

Simulations of mutant SecM were performed with a force loading rate of 50 pN/ns. 20 independent trajectories were generated for each mutant.

Simulations of VemP were performed analogously to SecM, with a force loading rate of 35 pN/ns. Initially, 30 simulations were run until the minimum of the pulling harmonic potential was 40 Å away from the initial position of the VemP N-terminal alpha carbon, corresponding to a simulation time of 80 ns. However, this was insufficient to disrupt the upper helix of VemP and further extension would bring the N-terminus of VemP near the periodic boundary of the simulation. To overcome this, after each trajectory was pulled for 80 ns, the simulation was halted and all amino acids of the nascent chain which were fully extended (i.e ϕ and ψ dihedral angles near 180 degrees) were removed. Simulation was then resumed, with the distance restraint adjusted such that the applied force at the end of the previous simulation and start of the new simulation were the same. This procedure was repeated until all VemP amino acids were fully extended. No more than two round of truncation and restart were required to fully extend the nascent chain.

### Trajectory analysis

#### Assigning states based on force profiles

To enable identification of states over the ensemble of trajectories, force profiles are compared after subtracting the linear rate of increase of force with time. Note that this rate of increase is not equivalent to the force loading rate when the spring constant of potential used to apply the pulling force is greater than the curvature of the energy landscape around a state [28], as was found to be the case in our simulations. To determine the observed rate of force increase, the beginning and end points of a linear segment are identified by finding all the local minima and maxima in a given profile. To avoid identifying spurious extrema caused by noise in the force profile, the profile is smoothed with a moving average filter and any extrema within 50 frames of another extremum are neglected, thus enforcing a minimum length for the identified linear segments (see Fig. 1C for example results). The slope of these segments can then be calculated, revealing a singly peaked distribution with fat tails (Fig. S1 A-C). The median of this distribution is then used as the slope to flatten each of the force profiles, which are subsequently binned and fit to a distribution composed of the sum of Gaussians to define the five states (Fig. S1 D-F). Each frame is then assigned to one of the five states, based on which of the five Gaussians has the highest probability density at the force associated with the frame.

#### Conformational analysis

After each frame has been assigned to a state based on the force profile, the conformational changes between states are characterized in terms of the dihedral angles. Specifically, for a given state *i* and amino acid *a*, the dihedral distance is defined by 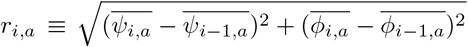. Averages of a dihedral angle are taken over all frames belonging to a given state and are calculated as circular quantities, i.e. angles are converted to Cartesian coordinates, averaged, and then converted back to an angle.

#### Calculation of tension and work

To calculate the tension on the nascent chain, the forces induced by extension or contraction of the backbone covalent bonds are recorded. In particular, for each amino acid the backbone bond distances between the amide nitrogen to alpha carbon, between the alpha carbon to carbonyl carbon, and between the carbonyl carbon and the subsequent amino acid’s amide nitrogen are recorded. These distances were then converted into forces based on the AMBER99SB-ILDN forcefield parameters [38]. The forces from these three bonds are then average into a single quantity for each amino acid. This calculation is performed overall all the frames in each state, as well as the equilibration trajectory in which no pulling force was applied. Tension on each amino acid is compared relative to the tension during equilibration and reported in Fig. 4C and Fig. S5.

The work to reach a state is calculated as the integral of the applied force from the start of a trajectory until all the dihedral conformational changes characteristic of the given state are observed. Specifically, state 2 is reached when Ser151 is straightened, state 3 is reached when Ser157, Gln158, and Ala159 are all straightened, and state 4 when Arg163 is straightened. A dihedral angle is straightened when the ψ dihedral angle is within 30 degrees of straight (180°) and remains so for 10 frames.

## 1 Acknowledgments

Anton 2 computer time was provided by the Pittsburgh Supercomputing Center (PSC) through Grant R01GM116961 from the National Institutes of Health. The Anton 2 machine at PSC was generously made available by D.E. Shaw Research. This work was also supported by a grant from NIGMS, National Institutes of Health, (R01GM125063) to TFM and MZ.

