## Supplemental Figures for "Force transduction creates long-ranged coupling in ribosomes stalled by arrest peptides"

### Supplementary Materials

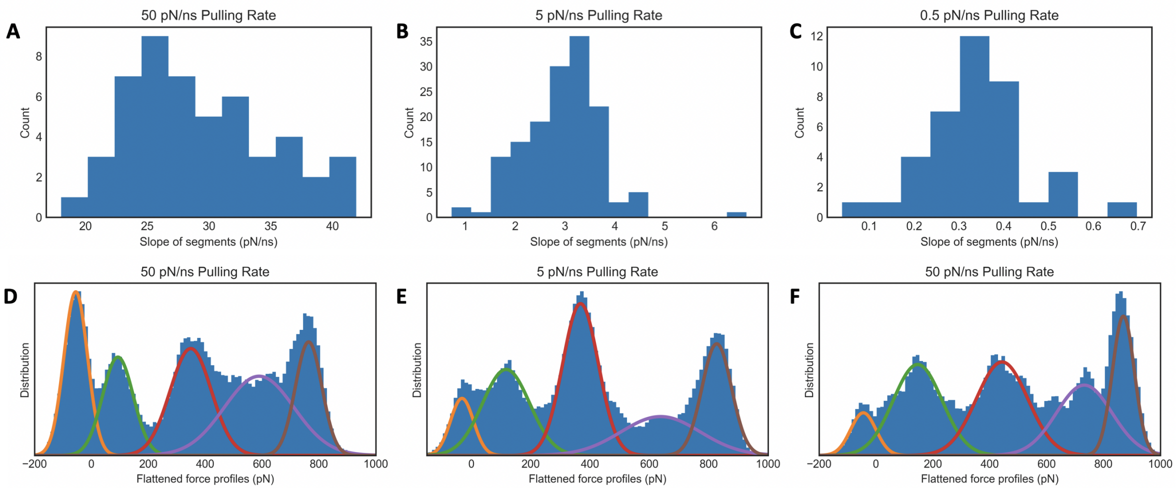


Figure 1: Identifying states based on force profiles. A) For all 30 50 pN/ns trajectory, linear segments were identified as described in the Methods. The distribution of the slopes of these segments is plotted. B) Same as A, but for all 30 5 pN/ns trajectories. C) Same as A, but for all 5 0.5 pN/ns trajectories. D) The median of the slopes of the linear segments plotted in A were subtracted from each force profile, generating flattened profiles as in Fig. 2A. The distribution of flattened force profiles is plotted. Superimposed is the results of fitting the distribution to the sum of five Gaussians. Each Gaussian is plotted separately. E) Same as D, but for 5 pN/ns trajectories. This panel is identical to Fig. 2B F) Same as D, but for 0.5 pN/ns trajectories.

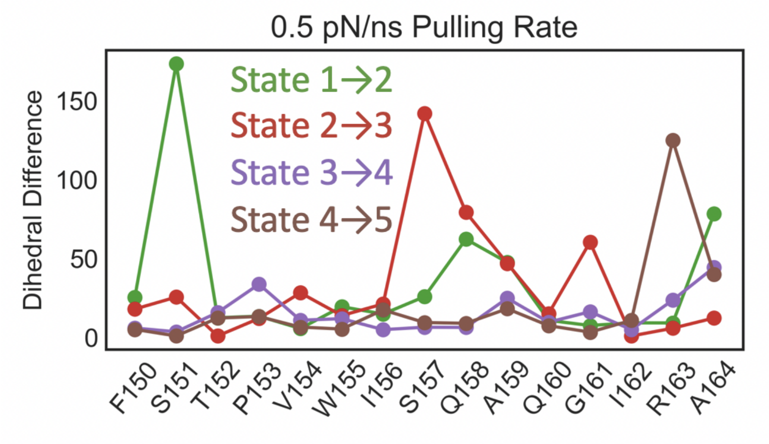


Figure 2: Conformational changes are quantified for each amino acid in terms of the distance between the $\phi$ and $\psi$ dihedral angles of two states. Data are from 0.5 pN/ns force loading trajectories. Data from 5 and 50 pN/ns trajectories are plotted in Fig. 3

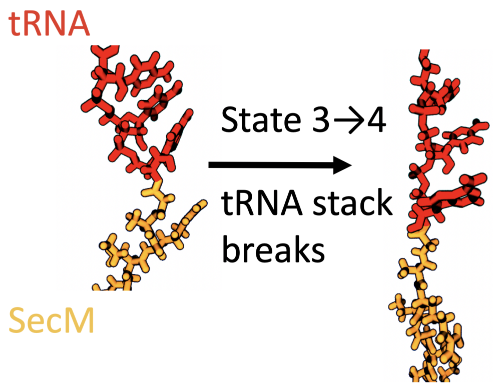


Figure 3: Representative conformations of the nascent chain and tRNA are displayed for states 3 and 4. Moving between the states requires significant disruption of the base-stacking interactions in the tRNA.

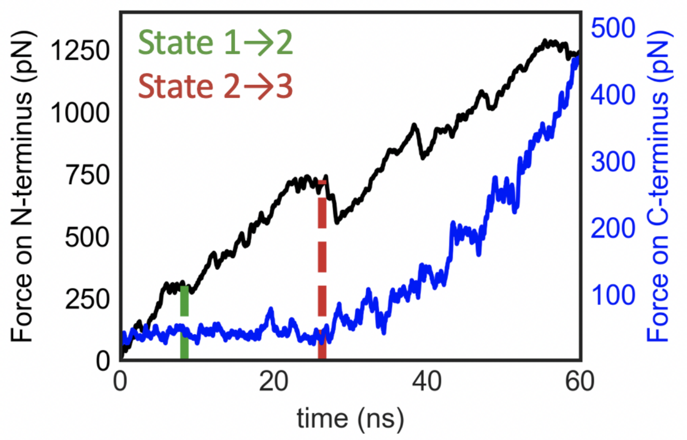


Figure 4: Simulations of SecM were run with the force on the N-terminus ramped at 50 pN/ns, but now with the C-terminal Gly165 fixed by a harmonic restraint. The applied pulling force on the N-terminus is plotted in black and the amount of force that is acting on Gly165 is plotted in blue. The figure is equivalent to Fig. 3B, except Gly165 was fixed and used to measure forces at the C-terminus rather than Arg163. Regardless of the precise residue in the C-terminus at which force is measured, no forces above the background are observed until state 3 is entered.

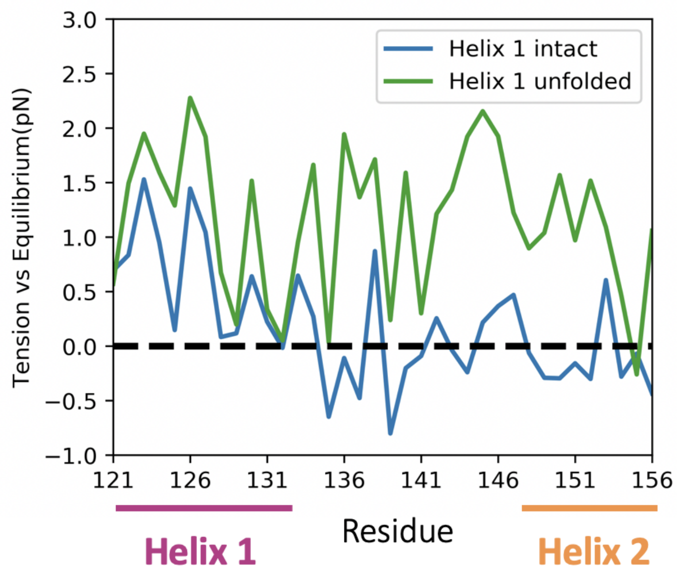


Figure 5: The tension along the VemP backbone relative to the tension during equilibration is averaged over all frames with helix 1 either intact or unfolded, then plotted per amino acid. The unfolding of helix 1 is measured by the straightness of Thr135 at the C-terminus of helix 1. When the dihedral angle is within 30 degrees of straight for more than 3 ns, the helix is defined as unfolded. While helix 1 is intact, the pulling force does not induce additional tension in helix 2, implying the propagation of force up nascent chain is inhibited. Only once helix 1 is unfolded does helix 2 experience increased tension.

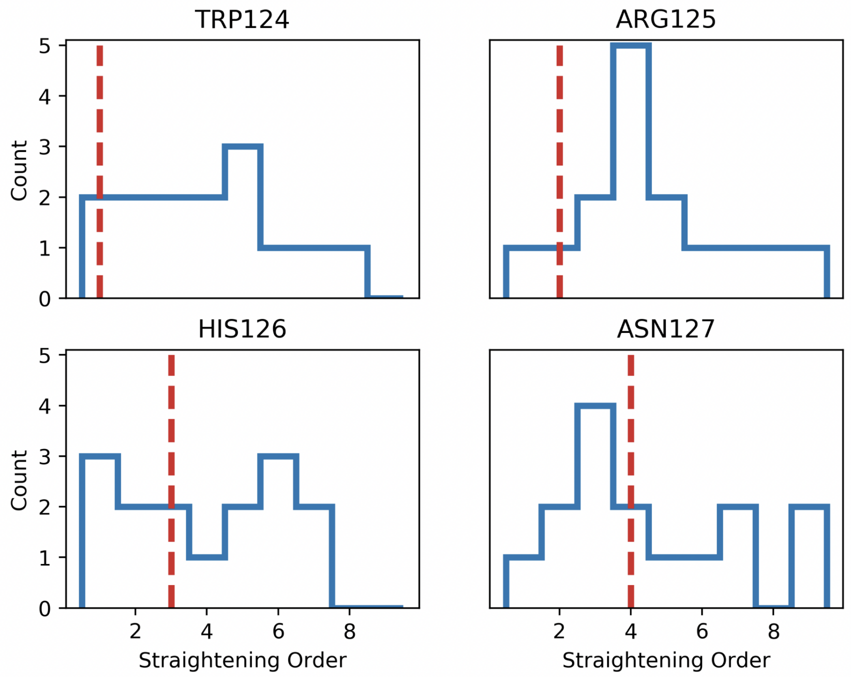


Figure 6: The order in which the dihedral angles of the first four amino acids in VemP helix 1 straightens is determined for each trajectory and the distribution of these orders is plotted. The expected order if the amino acids straightened consecutively is indicated with the red dashed line. For example, Arg125 is most often the 4th amino acid in helix 1 to straighten, despite being the 2nd amino acid from the helix N-terminus. All four dihedral angles show significant heterogeneity in the order of straightening across different trajectories.

#### Estimation of energy landscape

Given a number of repeated and stochastic trajectories of force-induced disruption of a bond, it is possible to estimate the free energy landscape of that bond. This estimation is typically performed by assuming one-dimensional double-well energy landscape that the system explores through thermal fluctuations, just as in Kramer's theory of chemical reaction rates[1]. The applied force distorts the underlying free energy landscape of the bond, acting to reduce the height of the energy barrier and increase the rate of crossing. Bond rupture occurs when the free energy barrier is crossed.

Numerous theories have been developed that provide strategies to estimate an energy landscape given different pulling protocols and force regimes [2, 3, 4]. In this work, we use the framework developed by Bullerjahn, Strum, and Kroy [5] which has the advantage of being directly applicable to the force loading protocol used in our molecular dynamics simulations, namely the application of external force via a stiff and rapidly moving harmonic potential. Their theory predicts the distribution of observed rupture forces given the parameters describing the energy landscape and the pulling protocol, i.e. $p(F|x^{\ddagger}, \Delta G^{\ddagger}, D; k_s v)$. $x^{\ddagger}$ is the distance from the minimum of the bound state to the location of the barrier maximum, $\Delta G$ is the height of the free energy barrier, $D$ is the diffusion constant, and $k_s v$ is the force loading rate used in simulation. $k_s v$ is chosen when setting up the simulation, while the remaining three parameters need to be estimated. This is done by finding the value of the parameters that maximizes $p(F|x^{\ddagger}, \Delta G^{\ddagger}, D; k_s v)$. The functional form for this probability is provided in [5], and assumes the underlying free energy landscape to be harmonic with a cusp at the barrier maximum.

Optimizing this probability to find $x^{\ddagger}$, $\Delta G^{\ddagger}$, and $D$ simultaneously produced unphysical results with estimates of $\Delta G^{\ddagger}$ and $D$ both orders of magnitude higher than expected for all transitions. The poor estimates are likely due to insufficient samples of stall-breaking events. Instead of performing a fully flexible fit, we first estimate the spring constant of the underlying harmonic energy landscape $k_l$, which fixes the ratio of $x^{\ddagger 2}$ to $\Delta G^{\ddagger}$ as $k_l = \frac{2\Delta G^{\ddagger}}{x^{\ddagger 2}}$. The estimate for the spring constant of the underlying landscape is obtained by first expressing the effective potential as the sum of the underlying landscape plus the moving harmonic trap, i.e.

$$U(x,t) = k_l x^2 + k_s (x - v_s t)^2$$

where $k_s$ is the spring constant of the applied harmonic potential and $v_s$ is the speed at which the applied potential is moved. Because the harmonic trap is moving with time, the energy minimum of the effective potential moves as well, specifically

$$x_{min}(t) = \frac{k_s v_s t}{k_s + k_l}$$

As the system tracks the moving minimum of the effective potential, the expected value of the applied force at a given time varies as:

$$\mathbb{E}[F(t)] = k_s \left(vt - x_{min}(t)\right) = k_s vt \left(1 - \frac{k_s}{k_s + k_l}\right)$$

The applied force is an experimental observable, and all the other parameters besides $k_l$ are known. This means that the spring constant of the underlying constant can be simply estimated from the slope of the applied force as a function of time:

$$k_l = \frac{k_s}{1 - \dot{F}/k_s v}$$

where $\dot{F}$ is the slope of the applied force. The estimation of $\dot{F}$ is described in Methods and results are shown in Fig. S1. The estimate of $k_l$ is informative because the spring constants of the underlying landscape and the applied potential are of a similar scale. This is not typically the case for optical tweezer experiments, for which $k_l \gg k_s$ and therefore $\mathbb{E}[F(t)] \approx k_s vt$, which has no $k_l$ dependence.

Estimating $k_l$ removes one free parameter from the optimization problem, as $x^\ddagger$ and $\Delta G^\ddagger$ are linked through $k_l = \frac{2\Delta G^\ddagger}{x^{\ddagger 2}}$. Still, fitting the observed forces at rupture to $p(F|x^\ddagger, \Delta G^\ddagger, D; k_s v)$ produced unphysical parameters. To further reduce the complexity of the fit, $x^\ddagger$ is fit to the median of the rupture forces while $D$ is held constant. By repeating this fit over each of the three pulling forces and finding the value of $D$ which leads to the best agreement between predictions of $x^\ddagger$, both $x^\ddagger$ and $D$ can be estimated (Fig. S7).

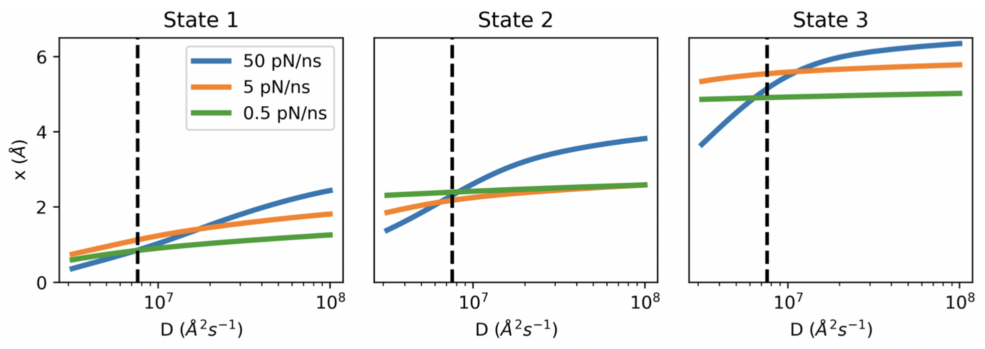


Figure 7: Each plot shows the optimum value for $x^\ddagger$ given a fixed value for the diffusion constant. Each plot corresponds to the barrier for exiting a given state, and the fits are performed separately for trajectories of different force ramping rates. The dashed lines indicate the value of the diffusion constant that produces the best agreement between estimates of $x^\ddagger$ across all three pulling speeds and states.

The above methodology allows fitting of a single-barrier landscape, however the the pathway of force-induced SecM restart clearly exhibits multiple, sequential barriers. To fit barriers beyond the first, it is necessary to find the location of the minimum of the state before the barrier of interest. This is done by assuming that the distance from the previous minimum to the barrier is the same as the distance from the barrier to the subsequent minimum (Fig. S 8). This significant and simplistic assumption is necessary because our trajectories do not exhibit the rebinding events that would be necessary to gain more detailed information on the shape of the landscape between a barrier and the subsequent minimum. Despite the simplicity of the assumption, we

Table 1: Optimal parameters that describe that energy barrier for leaving states 1-3. Confidence intervals were determined by bootstrapping over trajectories.

| | $x^{\ddagger}(\AA)$ | $x^{\ddagger}$ 90% CI | $\Delta G^{\ddagger}(k_BT)$ | $\Delta G^{\ddagger}$ 90% CI | $k_0(s^{-1})$ | $k_0$ 90% CI |
|---|---|---|---|---|---|---|
| State 1 | 1.0 | 0.83 | 4.2 | 3.3 | $1.0 \times 10^6$ | $6.3 \times 10^4$ |
| | | 1.3 | | 8.1 | | $2.3 \times 10^6$ |
| State 2 | 2.5 | 2.2 | 27 | 21 | $4.3 \times 10^{-4}$ | $6.7 \times 10^{-25}$ |
| | | 3.9 | | 76 | | $8.9 \times 10^{-2}$ |
| State 3 | 5.3 | 5.0 | 120 | 110 | $2.8 \times 10^{-45}$ | $2.7 \times 10^{-61}$ |
| | | 6.0 | | 160 | | $1.6 \times 10^{-40}$ |

do not observe any states rupturing before their predicted minima, which would be clear indicator of a poor choice of landscape. Having established a method to identify the minima of a state given the parameters estimated for the previous state, we can iteratively estimate the landscape of each state (Fig. S7, Table S1).

The estimated parameters clearly indicate that the barrier between state 1 and 2 cannot be the sole barrier before restart of translation, as it can be overcome by thermal motion with a rate of over $10^4\ s^{-1}$. The crossing of the barrier between states 3 and 4 cannot be required for restart of translation because the crossing time is far too slow at physiological forces, e.g. Bell's law predicts that with a force of 10 pN the barrier will be crossed with a rate $k_0 e^{Fx^{\ddagger}/k_BT} = 10^{-44} s^{-1}$. Note that both barriers are still implausibly relevant even at the ends of the 90% confidence intervals.

By contrast, the barrier between state 2 to 3 has a more physiologically sensible rate constant of $4.3 \times 10^{-4}$ which drops to $5.6 \times 10^{-3}$ with a 10 pN force, assuming Bell's law. Although the confidence intervals on these estimates are wide, this barrier appears to be substantial yet still physiologically surmountable. In combination with the fact that crossing this barrier allows the force to affect the conformation at the C-terminus of SecM, the barrier between state 2 and 3 is likely to be the key barrier to force induced restart of translation.

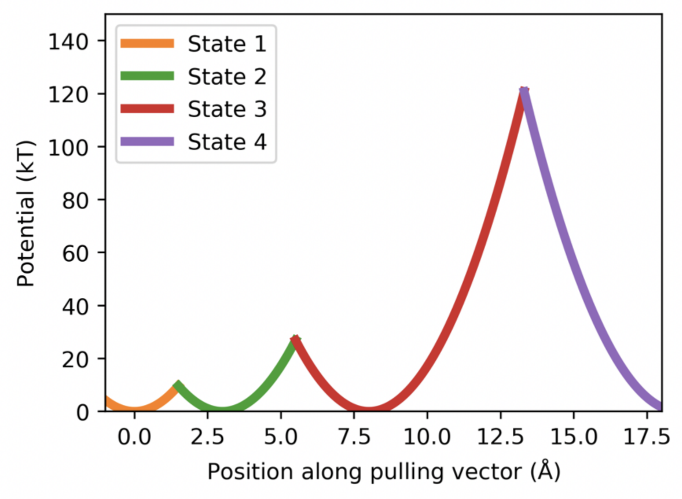


Figure 8: Estimated potential energy landscape for force-induced SecM restart. Each potential energy well is colored by the state it corresponds to.

##### SI Movie 1

A representative trajectory is shown with a pulling rate of 5 pN/ns and a total time of 600 ns. The ribosome is colored in blue, with the fully flexible region colored in dark blue and the restrained outer shell in cyan. The nascent chain is colored in orange, and the tRNA in red. Ribosomal atoms nearer to the viewer than the nascent chain are removed. As portions of the nascent chain straighten in response to the applied force, the fully straightened portions are colored in white. This region grows from the N-terminus to the C-terminus in distinct steps, corresponding to the five states characterized in Figs. 2 and 3.
